# The mitotic stopwatch limits the proliferation of whole genome doubled cells

**DOI:** 10.1101/2025.09.18.676992

**Authors:** LA Allan, R Foy, M Keli, AT Saurin

## Abstract

Whole genome doubling (WGD) is a frequent event in tumourigenesis that promotes chromosomal instability and tumour evolution. WGD is sensed indirectly due to the presence of extra centrosomes, which activate the PIDDosome to induce a p53-dependent G1-arrest. Here we uncouple WGD from centrosome amplification and show that p53 still arrests tetraploid cells, but via the mitotic stopwatch; a p53/53BP1/USP28-dependent pathway that causes G1-arrest following an extended mitotic delay. Mitotic timing is unaffected by WGD, but the threshold mitotic delay needed to invoke a G1-arrest is reduced. This sensitivity to the stopwatch mechanism is not associated with altered levels of mitotic stopwatch components, but instead, is associated with enhanced p21 concentrations prior to mitosis. Similar effects are observed in diploid cells treated with CDK4/6 inhibitors to double their size, implicating G1 delays and cell size as key determinants of stopwatch sensitivity. This ability of the mitotic stopwatch to arrest proliferation after increases to genome or cell size has important implications for the progression and treatment of cancer. It also demonstrates that the stopwatch pathway can sense more than just mitotic delays.

## Introduction

Whole-genome doubling (WGD) occurs in almost 40% of human tumours at some stage in their evolutionary history, making it one of the most common genetic alterations in cancer^1–5^. WGD is associated with chromosomal instability, rapid tumour evolution and poor patient prognosis^1,3,4,6,7^. This reinforces the need to understand the mechanisms that prevent proliferation after WGD, and to understand why these fail to do this effectively in cancer cells.

It is well-established that the tumour suppressor p53 protects cells against WGD by arresting tetraploid cells in G1-phase of the cell cycle^8–10^. This explains why TP53 mutation is strongly associated with WGD in human tumours, and why, when these events co-occur, p53 mutation almost always precedes WGD^1–3^. Mechanistically, p53 is not thought to sense increased ploidy directly, but rather, it senses the presence of extra centrosomes which are amplified together with chromosomes during S-phase. These extra centrosomes activate the Hippo pathway^11^ or the PIDDosome complex^12–14^ to inhibit or cleave MDM2, respectively, thus stabilising p53 and inducing a p21-mediated cell cycle arrest. While this provides a mechanism to restrain the proliferation of cells after WGD, it remains unclear whether genome amplification can also be sensed directly by p53 or other pathways independently of centrosomes. This is important because without such a mechanism, genome amplified cells could easily evade detection if centrosomes are not co-amplified.

We address this question here by creating whole genome doubled cells with normal centrosome numbers. We find that p53 still inhibits the proliferation of these cells, but via the mitotic stopwatch (also known as the mitotic surveillance pathway): a p53/53BP1/USP28-dependent mechanism that arrests cells in G1 after mild mitotic delays^15,16^. WGD sensitizes cells to the stopwatch mechanism, and this sensitivity is associated with elevated p21 levels prior to mitosis; an effect that is also observed following cell cycle delays that lead to cellular enlargement. These data suggest that the mitotic stopwatch does not only detect mitotic delays, it can also respond to earlier cell cycle defects that lead to increased genome or cell size.

## Results

### p53 restricts the proliferation of whole genome doubled cells with normal centrosome numbers

To address whether cells could sense genome amplification in the absence of centrosome duplication, we generated tetraploid (4N) cells with normal centrosome numbers using a telomerase-immortalised RPE1 cell line containing auxin-degradable PLK4^17^. PLK4 is required for centrosome duplication in S-phase^18^, therefore by treating cells with auxin (IAA) prior to inducing cytokinesis failure we were able to generate tetraploid cells with normal centrosome numbers (Fig.1A-B). We readily generated 4N clones using this protocol in RPE1 cells in which p53 was knocked out (p53-KO) or knocked down (p53sh), but the numbers of clones generated in p53-proficient RPE1 cells was significantly reduced (Fig.1C and S1A-D). Moreover, we managed subsequently to generate 8N clones with normal centrosome numbers from these 4N cells, but only in the p53-KO or p53-sh backgrounds (Figure S1C-E). This implies that p53 acts as a barrier to genome amplification despite normal centrosome numbers. To assess this more directly, we utilised the p53sh cells in which endogenous p53 can be re-expressed upon removal of doxycycline (Fig S1F). This caused tetraploid p53sh cells to arrest but had little effect on diploid p53sh cells, demonstrating that p53 can restrain the proliferation of cells after WGD (Fig. 1D-F). Since this occurs in the presence of normal centrosome numbers, it suggests that p53 can more directly monitor increased ploidy. Similar results were obtained in 8N cells after reinstatement of p53 (Fig. S1G).

**Figure 1:**
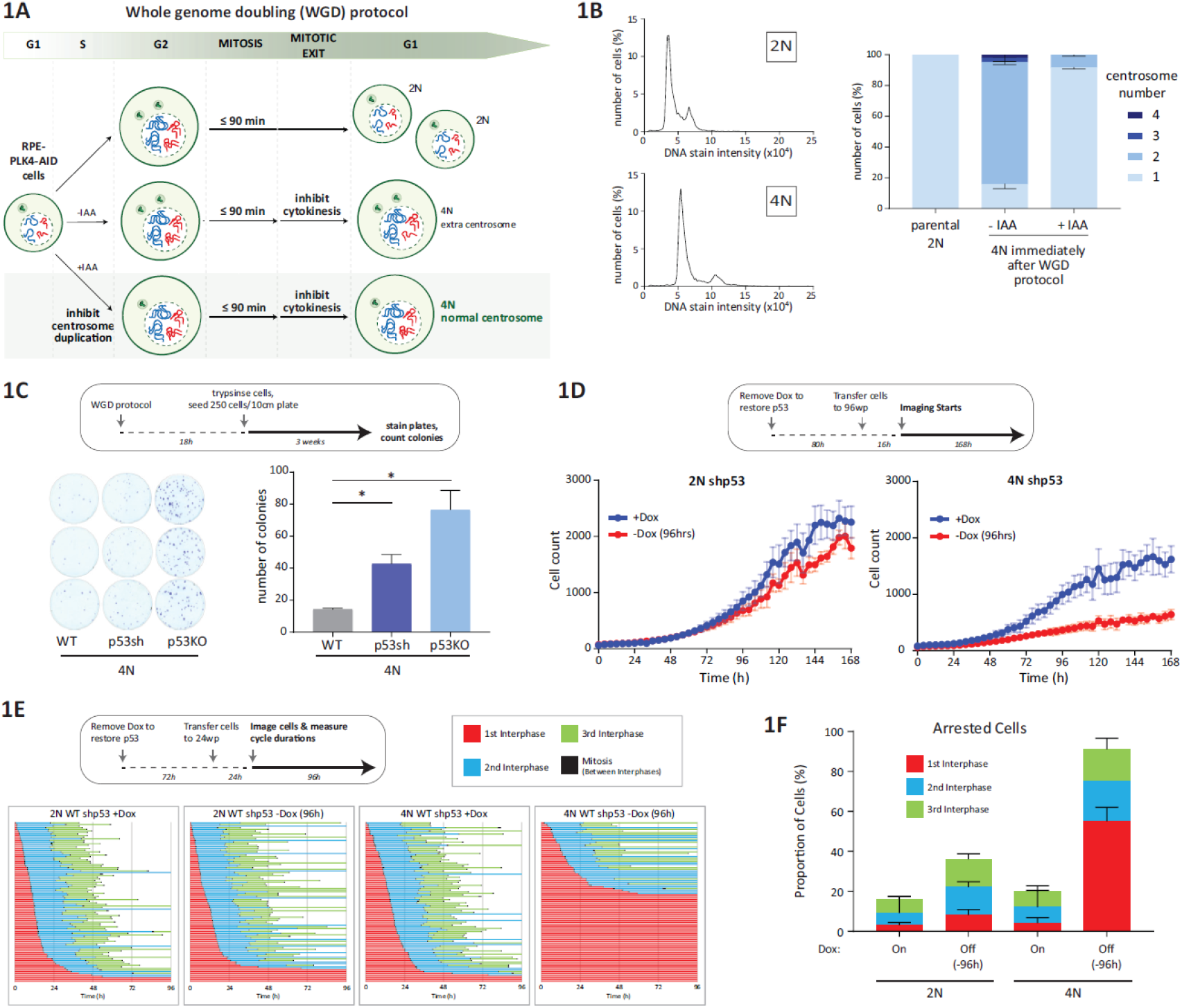
p53 limits proliferation in whole genome doubled cells with normal centrosome numbers. **A)** Scheme outlining the generation of whole genome doubled cells with normal centrosome numbers using auxin (IAA) degradable PLK4. **B)** Propidium Iodide staining of parental (2N) and WGD (4N) RPE1 PLK4-AID cells following the protocol described in A (left panel) and quantifications of centrosome numbers in these cells immediately following WGD protocol (right panel, 46-54 cells/experiment, n=3). Bars represent means ± SD. **C)** Colony forming assays of WGD cells following the protocol described in A. Following the WGD protocol a total of 250 cells were seeded at low density and allowed to form colonies for 3 weeks. Bars represent means ± SD, n = 3. \**P* < 0.05, unpaired two-tailed *t*-test. **D)** Line graphs representing cell counts of doxycycline-inducible p53-shRNA 2N/4N cells over 7 days either in the presence of doxycycline (blue) or starting 96 hours after doxycycline withdrawal (red). Data shown is the mean of 3-5 repeats +/-SEM. **E)** Single cell traces of doxycycline-inducible 2N/4N p53-shRNA cells (each line represents one cell lineage) showing three cell cycles in the presence of doxycycline or starting 96 hours after doxycycline withdrawal, as indicated. Fifty cells were analysed at random for each repeat and two experimental repeats are displayed (100 cells total). **F)** Quantification of observed cell cycle arrests from the data shown in E. Note, cell lineages which do not complete three cycles within the 96hr window are considered to be arrested. Bars show mean + SD from 2 repeats.

### The mitotic stopwatch limits cell proliferation after WGD

One mechanism by which p53 can arrest cell proliferation is via the mitotic stopwatch: a p53/53BP1/USP28-dependent pathway that is activated by mitotic delays to induce p21 and arrest cells in the subsequent G1^15,16^ (Fig. 2A). In diploid RPE1 cells, the mitotic stopwatch is activated when mitosis is delayed for over 90 minutes^19,20^, however, this threshold was decreased to approximately 60 minutes in three independent tetraploid RPE1 clones (Fig. 2B). Mitotic duration was not extended in these tetraploid cells, and this was generally lower than the 60 mins needed to invoke a full G1 arrest (Fig. 2C). However, multiple sub-threshold mitotic delays were recently shown to cumulatively activate the stopwatch to cause cell cycle exit^21^. Therefore, to test if the mitotic stopwatch helps to restrain the proliferation of tetraploid cells, we created USP28 knockouts in tetraploid RPE-PLK4 AID cells (Fig. S2A). We chose to target USP28, because the other central stopwatch component, 53BP1, is a mediator of the DNA damage response and this has also been implicated in p53 activation after WGD^22^. As expected, the stopwatch was abolished in both tetraploid clones (Fig. S2B). Next, we created USP28 knockouts in the tetraploid doxycycline-inducible p53-shRNA background (Fig. S2C). Figure 2D shows that removal of doxycycline caused proliferation to dramatically slow in parental tetraploid cells, but this was rescued in tetraploid USP28-KO cells. This effect was also observed in a separate USP28 knockout clone, (Fig. S2D). Single cell profiles from live cell movies demonstrates that this reduced proliferation was due to a large portion of cells arresting in the cell cycle within a few days after doxycycline removal (Fig. 2E-F and S2E). This demonstrates that reinstating p53 arrests the proliferation of tetraploid cells via a USP28-dependent pathway. Considering that USP28 is an essential component of the stopwatch^15,16^, and that the stopwatch threshold is reduced in tetraploid cells (Fig.2B), we conclude that the mitotic stopwatch acts as a barrier to proliferation after WGD.

**Figure 2:**
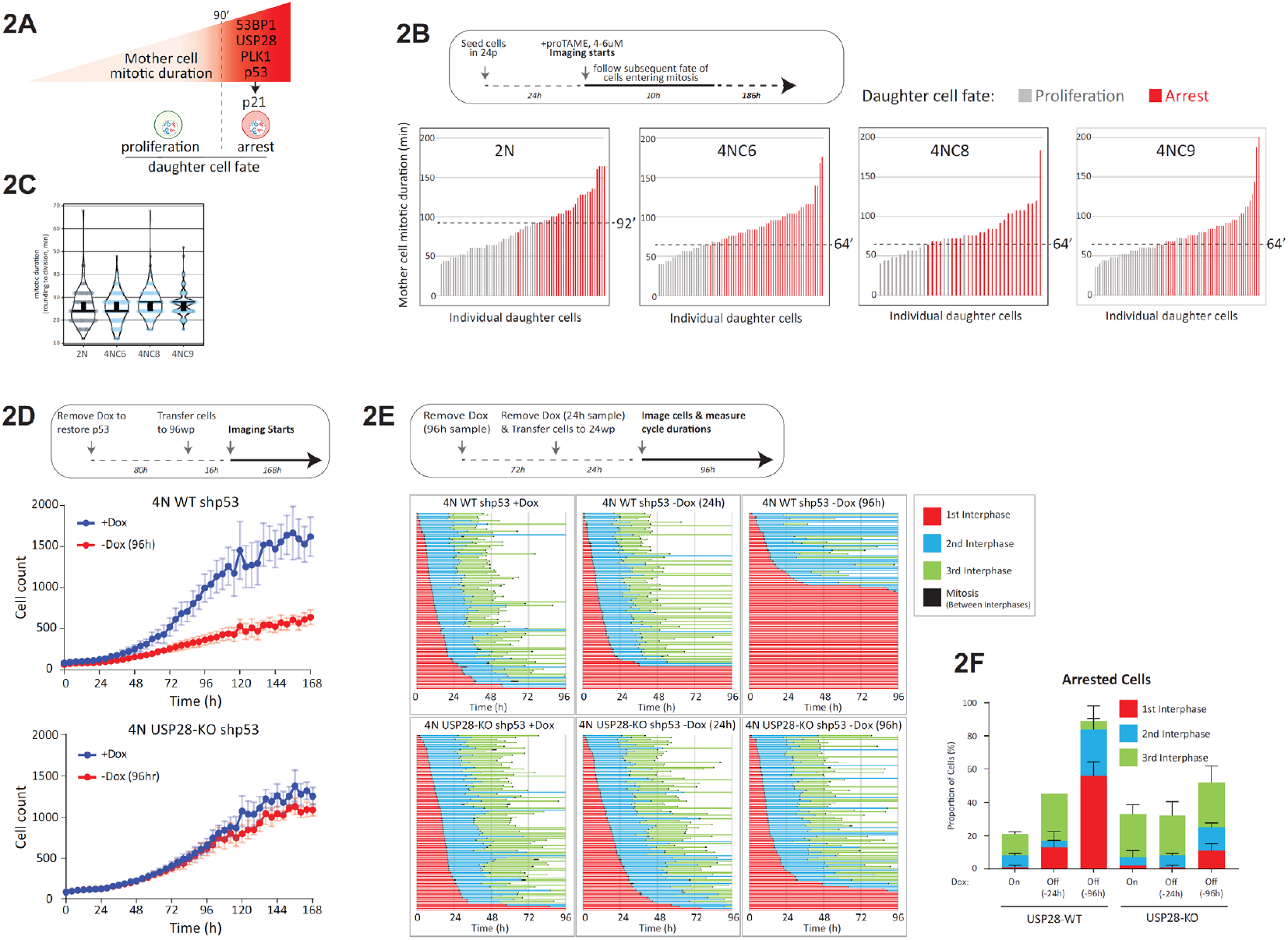
The mitotic stopwatch is a barrier to proliferation after WGD. **A)** Scheme to explain the mitotic stopwatch. **B)** Plots showing daughter cell fate as a function of mother cell mitotic duration in parental (2N) and three different WGD clones following treatment with proTAME to induce mild mitotic delays of varying lengths. Only cells entering mitosis in the first 10h were followed for daughter cell fate and cells which fail to re-enter mitosis within the remaining 186 h window are considered to be arrested. Each bar represents the time spent in mitosis for a single mother cell while the colour of the bar represents daughter cell fate (grey = proliferation, red = arrest). Data from 2 independent experiments. **C)** Violin plots representing unperturbed mitotic durations of parental RPE PLK4-AID 2N cells and 4N clones. Data are from from 3 experiments. The horizontal line represents the median time in mitosis while the thick vertical line gives the 95% CI. **D)** Line graphs representing cell counts of USP28-WT or KO doxycycline-inducible p53-shRNA cells cultured for 7 days either in the presence of doxycycline (blue) or starting 96 hours after doxycycline withdrawal (red). Note, the data displayed for the 4N WT is the same as that shown in figure 1D, with experiments performed at the same time. Data shown is the mean of 3-5 repeats +/-SEM. **E)** Single cell traces of 4N USP28-WT (above) or -KO (below) doxycycline-inducible p53-shRNA cells (each line represents one cell lineage). Traces show three cycles in cells cultured in the presence of doxycycline (left panels) or following the removal of dox for 24 or 96h, as indicated. Fifty cells were analysed at random for each repeat and two experimental repeats are displayed). **E)** Quantifications of cell cycle arrest in D. Note, cell lineages which do not complete three cycles within the 96hr window are considered to be arrested. Bars show mean + SD from 2 repeats.

### The mitotic stopwatch also limits cell proliferation after cell enlargement

Two major differences between tetraploid and diploid cells are increased DNA content and increased cell size. We hypothesised that the increased DNA content could reduce the stopwatch threshold, for example, by doubling kinetochores which have been implicated in the stopwatch mechanism^23–25^. To test this hypothesis, we created similarly sized diploid cells by treating with the CDK4/6 inhibitor, palbociclib (CDK4/6i). When diploid RPE1 cells are arrested with CDK4/6i they continue to grow in size during G1^26,27^, reaching the size of tetraploid cells after 1-2 days of arrest (Fig. 3A). Interestingly, these enlarged cells also displayed a reduced stopwatch threshold when released from that G1 arrest, with relatively mild mitotic delays of between 60 and 90 minutes invoking a penetrant cell cycle arrest (Fig. 3B). This demonstrates that prior cell cycle delays can also modify the stopwatch threshold, implicating cell enlargement during these delays as a possible mechanism. To test if cell enlargement was crucial, we arrested cells in CDK4/6i but restricted cell growth by inhibiting the mTOR pathway, as previously^26,27^. This prevented cell enlargement (Fig. 3A) and normalised the mitotic stopwatch threshold back to 90 mins (Fig. 3B), demonstrating the cell growth during a cell cycle delay can modify the stopwatch threshold afterwards.

**Figure 3:**
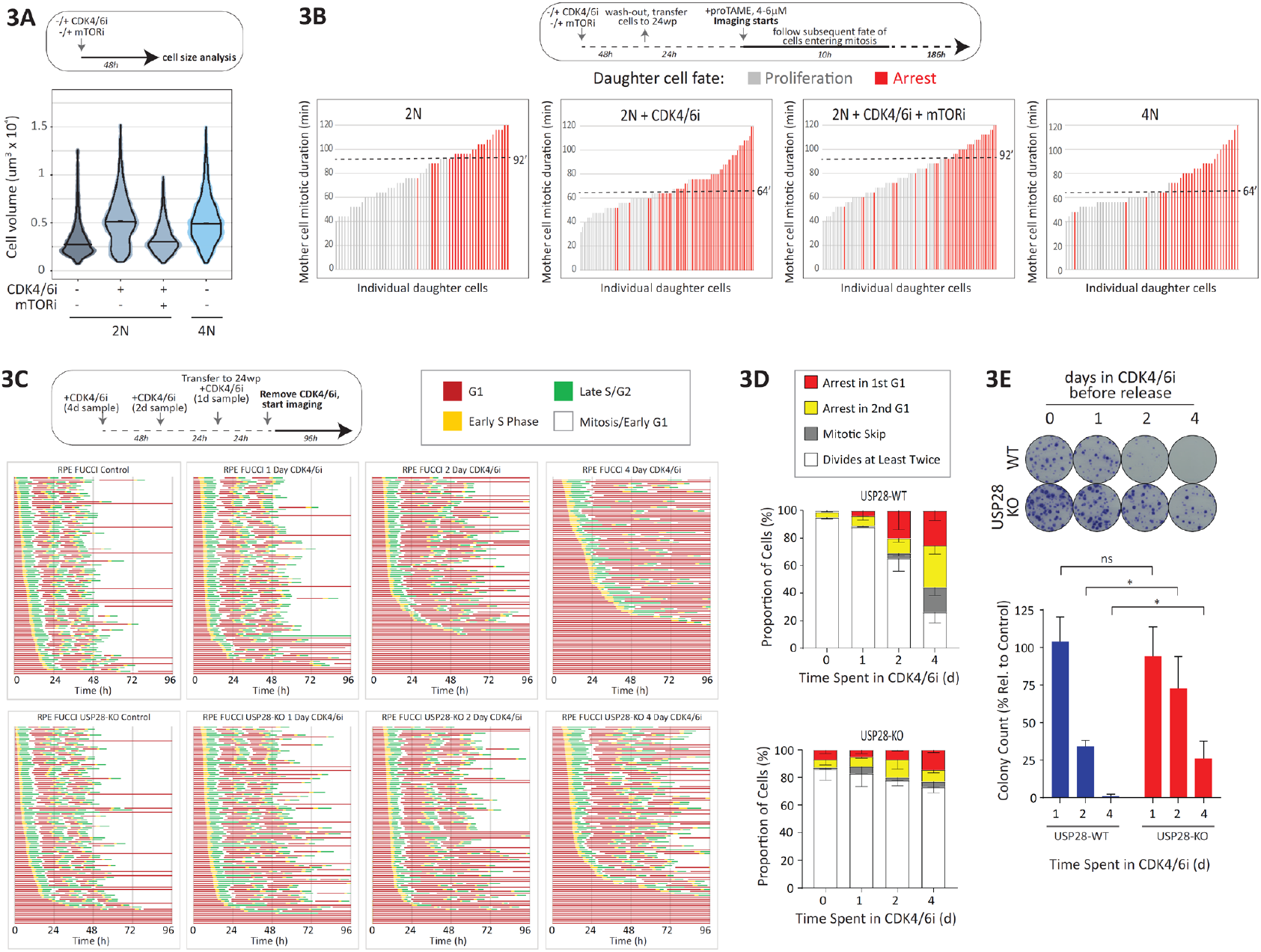
The mitotic stopwatch is a barrier to proliferation after cell enlargement. **A)** Violin plots showing cell volumes of parental RPE1 PLK4-AID cells before and after 2 days of CDK4/6 inhibitor (palbociclib) -/+ mTORi treatment compared to WGD (4N) cells. The horizontal line depicts the median and the thick vertical line represents the 95% CI, n=4. **B)** Plots showing (cont.) *(Continued from previous page)* daughter cell fate following mother cell mitotic duration in parental and WGD RPE1 PLK4-AID cells following treatment with proTAME to produce mild mitotic delays of varying lengths. Parental cells were either left unperturbed or first arrested in G1 with a CDK4/6 inhibitor (palbociclib) for 2 days either alone (to create large cells) or in combination with an mTOR inhibitor (to keep arrested cells small) before being released from G1 for 24hrs prior to the addition of proTAME. Only cells entering mitosis in the first 10h were followed for daughter cell fate and cells which fail to re-enter mitosis within the remaining 186 h window are considered to be arrested. Each bar represents the time each mother cell spends in mitosis with the colour of the bar showing daughter cell fate with grey representing continued proliferation and red depicting cycle arrest. Data are from >60 cells, n=2. **C)** Cell cycle profiles of individual USP28-WT (above) and -KO (below) RPE1-FUCCI cells (each bar represents one cell lineage) which were either asynchronous or treated with a CDK4/6 inhibitor (palbociclib) for 1, 2, or 4 days. Cell imaging was started immediately after removal of the inhibitor. Fifty cells in G1 during the first frame of the movie were selected at random then tracked and analysed for 3 divisions or until the movie ended. Data from 2 repeats is shown to give 100 cells in total. **D)** Stacked bars quantifying the proportion of defects observed in the first two cell cycles of the traces shown in C. Each bar represents the mean data from 2 repeats -SEM. **E)** Representative images and quantifications of colony forming assays in USP28-WT and -KO RPE

Cell enlargement after CDK4/6 inhibition causes cells to enter senescence in a p53-dependent manner^26–28^. We hypothesized that the mitotic stopwatch could contribute to this senescence entry if enlarged cells reach mitosis after CDK4/6 inhibitor washout. In agreement, the G1-arrest observed in enlarged cells that continue to cycle after washout from CDK4/6i is partially rescued in USP28 knockout cells (Fig. 3C-D), which is also sufficient to reduce long-term cell cycle arrest (Fig. 3E). Therefore, although cell enlargement following G1 delays can drive p53-dependent senescence directly from G1 or the subsequent G2^26–29^, if these cells manage to reach mitosis they can exit the cell cycle in the next G1 via a mitotic stopwatch-dependent mechanism.

### Sensitivity to the mitotic stopwatch is associated with elevated p21 in G2 cells

We next focussed on characterising molecularly why tetraploid or enlarged cells were particularly sensitive to the mitotic stopwatch. This could be due to stronger or quicker activation of the stopwatch mechanism during mitosis, or alternatively, it may represent enhanced sensitivity to its downstream effects during G1. One possibility is that the balance of mitotic stopwatch components, such as 53BP1, USP28, could be altered to affect the kinetics of stopwatch activation during mitosis^15,16^. Alternatively, factors needed to assemble this complex during mitosis, such as PLK1^21,24^, could also be affected. However, analysis of protein concentrations during G2 or mitosis, demonstrates that the levels of these components are unaffected by genome size (Fig. 4A) or cell size (Fig. 4B). 53BP1 is recruited to kinetochores and then released in a PLK1-dependent manner, which may affect the stopwatch response^24,25^. However, the kinetochore levels of 53BP1 or PLK1 were unaffected by genome size or cell size (Fig. 4A-B and S3). Mitotic degradation of the p53 ubiquitin ligase, MDM2, has been proposed to sense mitotic delays and therefore impact on the mitotic stopwatch response^30,31^. However, MDM2 levels during mitosis were unaffected after WGD and even elevated slightly by increased cell size, which is predicted to strengthen rather than sensitize the stopwatch (Figs. 4A-B).

**Figure 4:**
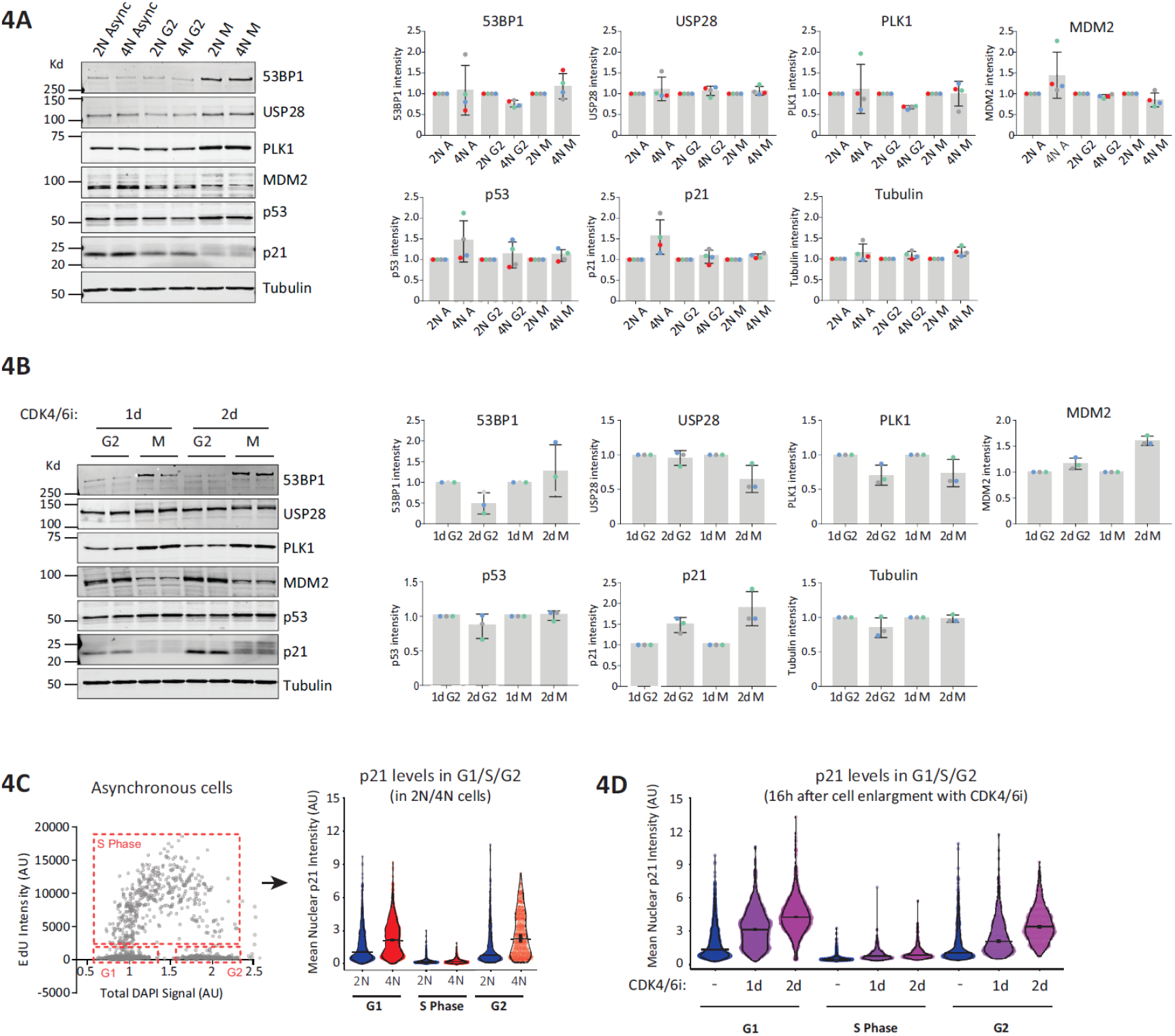
p21 is elevated following WGD or cell enlargement. **A)** Representative images (left) and associated densiometric quantifications (right) of western blots with lysates from RPE1 PLK4-AID parental (2N) and WGD (4N) cells when asynchronous or in either G2 or mitosis. Cells were first arrested in G1 with a CDK4/6 inhibitor (palbociclib) at high confluence to synchronize cells without causing aberrant cell growth. After 24hrs, cells were released from G1 and into STLC to prevent cells completing mitosis. After a further 16-17hrs all mitotic cells arrested in STLC were collected by shake-off and all remaining adherent cells were taken as the G2 samples. Quantifications shown are the mean +/-SD, n=4. **B)** Representative images (left) and associated densiometric quantifications (right) of western blots with lysates from RPE1 PLK4-AID parental cells following treatment with a CDK4/6 inhibitor (palbociclib) for either 1 or 2 days prior to release into STLC for 16-17h to collect G2 and M samples, as in A. Quantifications shown are the mean +/-SD, n=3. **C)** Left: a representative image of the cell cycle distribution produced by pulsing asynchronous RPE PLK4-AID cells with EdU for 30 minutes prior to fixation. EdU intensity was then plotted against total DAPI signal per cell allowing for cell cycle gating. Right: Violin plots depicting mean nuclear p21 intensity in asynchronous parental (2N) and WGD (4N) RPE1 PLK4-AID cells when in G1, S phase or G2. Horizontal bars represent the median from 3 repeats with the thick vertical bar showing the 95% CI. **D)** Quantification of nuclear p21 intensity in G1, S phase, and G2 in RPE1 PLK4-AID cells before and after 1-2 days of treatment with a CDK4/6 inhibitor (palbociclib). Note the data representing no palbociclib treatment is the same as the 2N data shown in C as these experiments were performed at the same time. Data displayed is from three repeats with horizontal lines representing the median and thick vertical lines showing +/-95% CI.

The output of the mitotic stopwatch is the production of p53 and p21, and the concentration of these proteins were consistently elevated in asynchronous tetraploid cells or in enlarged diploid cells as they progressed through G2 and mitosis (Figure 4A-B). The increased p21 in enlarged cells is unrelated to the mitotic stopwatch mechanism because these levels were elevated before cells transited through the first mitosis after CDK4/6i treatment. This suggests that the higher basal p21 prior to mitosis could sensitize cells to the additional p21 produced by the stopwatch. To test if p21 levels could also be elevated immediately prior to mitosis in asynchronous tetraploid cells, we performed p21 immunofluorescence in asynchronous cells and used DAPI and EdU staining to gate these cells into G1, S and G2 populations. Figure 4C shows that p21 concentrations are increased after WGD, and this occurs in both G1 and G2 cells. Note, the absence of p21 in S-phase was expected since p21 is specifically degraded by CRL4^Cdt2^ and SCF^Skp2^ ubiquitin ligases during DNA replication^32–35^. We performed the same assay 16h after treatment with CDK4/6i for 1 or 2 days to increase cell size, and we observed that p21 increased in both G1 and G2 cells in proportion to cell size (Fig. 4D). Note that these are mean nuclear p21 intensities, which provides an estimate of protein concentration within the nucleus. The increases are also consistent with those observed by Western (Figure 4B), which represents difference in total protein concentrations. In summary, increases to cell size and genome size elevates p21 concentration immediately prior to mitosis, which we hypothesise sensitizes cells to further elevation of p21 by the mitotic stopwatch.

If elevated basal p21 sensitizes tetraploid cells to the stopwatch mechanism, then reducing this basal p21 back to diploid levels should similarly reduce stopwatch sensitivity. CRISPR/Cas9 was used to decrease p21 copy number in tetraploid cells, thereby reducing p21 expression levels (Figure 5A). This was sufficient to decrease basal p21 in both G1 and G2 back to levels that approximate diploid RPE1 cells (Fig. 5B). This was also sufficient to increase the threshold mitotic timing needed to induce full G1 arrest back to 90 minutes, which was equivalent to the stopwatch threshold in diploid cells (Fig. 5C). Knockout of p21 completely abrogated the stopwatch, as expected (Figs. 5A-C). We conclude that WGD and cell enlargement sensitize cells to the mitotic stopwatch due to elevated basal p21. This helps to restrain the proliferation of these cells, which has significant implications for cancer progression after WGD and for cancer treatment with cell cycle inhibitors that increase cell size^36^.

**Figure 5:**
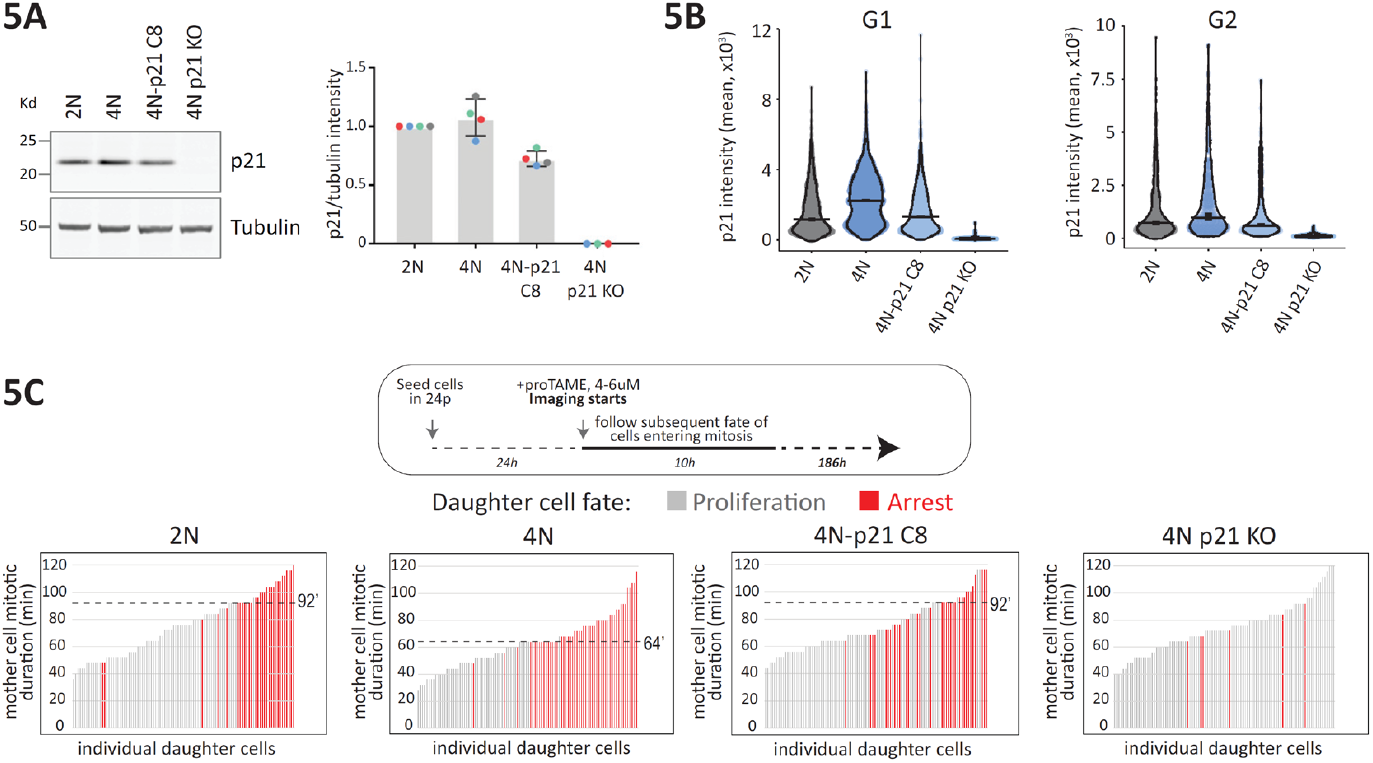
Partial depletion of p21 in WGD cells rescues stopwatch sensitivity. **A)** Representative images (left) and quantification (right) of western blots validating the partial reduction (4N-p21 C8) or complete loss (4N p21 KO) of p21 in WGD (4N) RPE1 PLK4-AID clones. Bars represent mean p21 intensity relative to tubulin -/+ SD, n=4. B) Violin plots quantifying nuclear p21 levels by immunofluorescence in parental (2N), WGD (4N), WGD p21-reduced (4N-p21 C8), and WGD p21-KO cells either in G1 (left panel) or G2 (right panel). Cell cycle segmentation was carried out as in 3C. Horizontal bar represents the median from 2 experimental repeats with the thick vertical line depicting the 95% CI. C) Plots showing daughter cell fate as a function of mother cell mitotic duration in parental (2N), WGD (4N), WGD p21-reduced (4N-p21 C8), and WGD p21-KO cells following treatment with proTAME to produce mild mitotic delays of varying lengths. Only cells entering mitosis in the first 10h were followed for daughter cell fate and cells which fail to re-enter mitosis within the remaining 186 h window are considered to be arrested. Each bar represents the time spent in mitosis for a single mother cell while the colour of the bar represents daughter cell fate with grey representing continued proliferation and red depicting cycle arrest. 50 cells per experiment, n=2.

## Discussion

The mitotic stopwatch pathway arrests cells following short mitotic delays, even if these delays result in normal chromosome segregations^15,31^. This is well-established to restrict proliferation of cells that have a wide range of mitotic defects^15^, but it is not immediately obvious why cells would evolve a mechanism to protect against mitotic delays that do not result in chromosome mis-segregations? Or to put this another way, why not just detect chromosome mis-segregations directly rather than mitotic delays that are not otherwise dangerous for cells? This manuscript goes some way to answering that question with the finding that the mitotic stopwatch does not only protect against mitotic delays or chromosome mis-segregations. Rather, it appears to protect against other abnormalities, such as delays in G1 progression and increases to cell size or genome size. In that sense it may monitor and protect cells against a wide range of cell cycle defects.

The role of the mitotic stopwatch in these situations appears to be to amplify pre-existing p53 signals to arrest cells in G1. It is possible that the mitotic stopwatch also acts to amplify other p53 signals to cause an arrest in G1 instead of G2. In agreement, cells that induce p21 mildly in G2 following DNA damage during S-phase, can arrest in the next G1 due to a spike in p21 after mitosis^37^. An independent study also supports the idea that the mitotic stopwatch helps to arrest cells with mild p53 elevation. CRISPR screening and live-cell imaging demonstrated that a range of genetic or drug perturbations that mildly induce p53 cause a proliferative arrest specifically in mitotic stopwatch-proficient cells (B. Mierzwa, A. Desai, K. Oegema, personal communication). It is unclear why the mitotic stopwatch amplifies p53 signals to arrest cells in G1, but one possibility is that this prevents WGD that may otherwise occur if cells arrest for prolonged periods in G2 and then skip mitosis^38,39^.

It will be important in future to determine the source of the enhanced basal p21 that helps to restrain cell proliferation after WGD. In response to increased cell size, p21 is induced by a combination and osmotic and replication stress^26,29^, which may explain why enlarged cells are sensitive to the stopwatch mechanism. Cells are also enlarged after WGD, but unlike cell enlargement with CDK4/6i, the cytoplasm:DNA ratio is better preserved after WGD. Considering that it is excess cytoplasmic growth and subsequent “genome dilution” that is implicated in the stresses experienced by enlarged cells^40^, we suspect that cell enlargement does not contribute to p21 induction after WGD. In fact, the opposite may even be the case, since decreased cytoplasm:DNA ratios were implicated in replication stress during the first S-phase after WGD^41^. Therefore, we speculate that the mitotic stopwatch helps to restrain proliferation of cells when the cytoplasm:DNA ratio becomes imbalanced, either due to changes in cell cycle duration, cell size or ploidy. Another possibility is that WGD changes the specific redox environment in cells, considering that oxidation of p21 has recently been linked to enhanced p21 degradation during G2 phase^42^.

These findings have important implications for cancer therapy, since CDK4/6 inhibitors are widely used to treat the most common subtype of metastatic breast cancer, but resistance is common^43^. Resistance is strongly associated with p53 mutation^44–46^, and this was recently linked to the inability of p53-deficient cells to enter senescence^46^. The loss of mitotic stopwatch activity could contribute to these effects, and potentially also to resistance against other chemotherapeutics that delay cells in G1 and increase cell size^36^. The mitotic stopwatch has previously been linked to senescence induction following chemotherapeutics agents that delay mitosis^21^, therefore this pathway may have evolved to protect against cell cycle delays in general. It is therefore important to carefully consider the mitotic stopwatch when assessing the response to any cell cycle inhibitors used in cancer therapy.

## Acknowledgements

We would like to thank the light microscopy facility at Dundee. We also thank Andrew Holland for the PLK4-AID RPE1 cell lines, Alexis Barr for comments on this manuscript, and Karen Oegema and Arshad Desai for sharing unpublished data. This work was supported by a Tenovus Scotland grant to A.T.S (T22-23), which funded R.F. and M.K., and a CRUK Programme Foundation Award (C4730/A21229) and Wellcome Investigator Award (222494/Z/21/Z) to A.T.S, which funded L.A.A.

## Author contributions

A.T.S. conceived the project, supervised the study and wrote the initial draft of the manuscript. L.A.A and R.F. generated all figures and edited the manuscript. L.A.A, R.F and M.K. performed all experiments and analysed all data.

## Materials AND Methods

### Cell culture and reagents

RPE-PLK4-AID cells^23^ were a gift from A.Holland, the RPE1-FUCCI cell line was published previously^39^ and HEK 293T cells were purchased from ATCC. Cells were authenticated by STR profiling (Eurofins) and screened to ensure a mycoplasma free culture every 4-8 weeks. Cells were cultured in DMEM (41966029, Thermo Fisher Scientific) supplemented with 10% FBS and 50 μg/ml penicillin/streptomycin (full growth media). The following drugs were used at the indicated concentrations throughout: Indole-3-Acetic Acid (0.1mM), Doxycycline (1μg/ml), STLC (10μM), Nocodazole (1μg/ml), Reversine (2μM), PF05212384 (30nM), Nutlin (5μM) and Digitonin (80ng/μl) were all purchased from Sigma Aldrich; PD0332991 (1.25μM) and puromycin (1μg/ml) from Santa Cruz Biotechnology; Centrinone (200ng/ml) and Etoposide (50μM) from MedChemExpress; proTAME (4-6μM) from R & D Systems; BI2536 (100nM) from Selleckbio and Zeocin (100μg/ml) from Invivogen.

### Plasmids and cloning

To generate a doxycycline-inducible p53-shRNA plasmid, we first subcloned a zeocin selection marker using BamHI-AatII and replaced the hygromycin selection marker in pEZ-Tet-pLKO-Hygro^47^ (a gift from Cindy Miranti - Addgene plasmid #85972; http://n2t.net/addgene:85972; RRID:Addgene_85972). Subsequently, oligos containing p53-shRNA sequences, (forward: 5’-CTAGCGACTCCAGTGGTAATCTACTTCAAGAGAGTAGATTACCACTGGAGTCTTTTTG-3’ and reverse: 5’-AATTCAAAAAGACTCCAGTGGTAATCTACTCTCTTGAAGTAGATTACCACTGGAGTCG-3’) were annealed and ligated into pEZ-Tet-pLKO-Zeo using NheI and EcoRI.

### Cell lines

RPE-PLK4-AID p53-KO cells were generated as described previously^29^. A similar protocol was used to knock out USP28 using a gRNA targeting exon 4 of USP28 (TGTAGCAACAGTGTCTTGAC). Transfected cells were selected for 2 weeks in centrinone (200ng/ml) before being plated at low density to generate single clones. Similarly, tetraploid cells with reduced p21 CDKN1A copy number and p21-KO cells were generated using a gRNA targeting exon 2 of p21 CDKN1A (CCGCGACTGTGATGCGCTAA). Transfected cells were selected in puromycin (10μg/ml) for 48 h, then in nutlin (5μM) for 24 h, before adding STLC (10μM) for a further 10 h. Mitotic cells, ie., those which had not arrested in nutlin and were predicted to be those in which p21 was fully knocked out, were removed and adherent cells incubated for a further 10 h in Nutlin + STLC. Again, mitotic cells were removed and remaining adherent cells were then replated at low density and allowed to recover in conditioned media for 3 weeks. Single clones were isolated and screened for reduced p21 protein levels or for complete knock-out by western blotting and immunofluorescence.

To generate doxycycline-inducible p53shRNA-expressing RPE-PLK4-AID and RPE-FUCCI cell lines, HEK-293T cells (ATCC) were transfected with pEZ-Tet-pLKO-Zeo-p53shRNA plasmid together with pRSV-Rev, pHCMVG and pMDLg/pRRE packaging vectors using LipofectAMINE 2000 (Thermo Fisher Scientific) according to the manufacturer’s instructions. After 24 h, the transfection mix was replaced with full growth media. Forty-eight hours after transfection, virus containing media was collected, filtered using a 0.45μM filter and diluted 1:3 with DMEM + 10% FBS. Cells were incubated with media-containing virus supplemented with polybrene (10μg/ml, Santa Cruz) for 24 h, followed by full growth media for a further 24 h. Transduced cells were then selected with Zeocin (100μg/ml) for 2 weeks.

### Whole genome doubling protocol

RPE-PLK4-AID cells at were incubated with Indole-3-Acetic Acid (IAA, 0.1mM) for 24 h. This was removed and cells were incubated with STLC (10μM) for 1 h before the addition of Nocodazole (1μg/ml) for 30 min. Mitotic cells were harvested by shake-off, washed in media containing nocodazole (1μg/ml), pelleted and replated in Nocodazole (1μg/ml), Reversine (2μM) and BI2536 (100nM) for 4 h to promote mitotic exit and cytokinesis failure and to allow cells to attach to the plate. Cells were then washed in full growth media every 30 min for 2 h and then cultured for a further 16h. Cells were then replated at low density (750 cells/15cm plate) in RPE-PLK4-AID conditioned media. Approximately 3 weeks later, individual colonies were isolated and analysed for ploidy status.

### Analysis of cell ploidy

Asynchronously growing cells were trypsinsed, resuspended in full growth medium containing digitonin (80μg/ml) and propidium iodide (PI, 50μg/ml, SIgma) and incubated at 37°C for 5 min. DNA content (PI intensity) was measured using a Chemometec NC-3000 Nucleocounter. Histograms of PI intensity were imported into Flowing Software version 5.2.1. Graphpad Prism 7 was then used to generate frequency distribution histograms.

### Cell volume analysis

RPE-PLK4-AID cells were plated at low density to prevent contact inhibition and incubated with CDK4/6 inhibitor, palbociclib (PD0332991, 1.25μM) -/+ mTOR inhibitor, PF05212384 (30nM) for 2 d. Cells were then trypsinsed, stained with acridine orange and DAPI (both Sigma), and cell diameter was measured using a Chemometec NC-3000 Nucleocounter. Histograms of cell diameter were imported into Flowing Software version 5.2.1 and cell diameter was then used to calculate volume using 4/3 πr3.

### Colony forming assays

250 cells were seeded in 10cm plates in triplicate and left to grow for 2 weeks. At the end of the assay, cells were washed twice in PBS and then fixed with 100% ethanol for 5 min. Plates were rinsed with water before being incubated with developing solution (1:1 ratio of 2% Borax:2% Toluene-D in water) for 5 min. Plates were then rinsed thoroughly with water and left to dry overnight. The plates were then scanned, and the number of colonies were quantified using ImageJ. This was performed by cropping to an individual plate and converting to a binary image. The fill holes, watershed and analyse particles functions were then used to count colonies. For colony forming assays after whole genome doubling, cells were allowed to recover in full growth medium for 16 h after going through the WGD protocol prior to seeding. To measure the effect of re-instating p53 expression on colony formation after whole genome doubling, RPE-PLK4-AID p53shRNA cells were cultured in the absence of doxycycline for 96 h prior to the WGD protocol.

### Immunofluorescence

Cells plated on High Precision 1.5H 12-mm coverslips (Marienfeld) were fixed with 4% paraformaldehyde (PFA) in PBS for 10 min or pre-extracted with 0.1% Triton X-100 in PEM (100mM PIPES, pH 6.8, 1mM MgCl2 and 5mM EGTA) for 1 min before addition of 4% PFA for 10 min. Pre-extraction was only performed for cells shown in Fig. S3. After fixation, coverslips were washed with PBS and blocked with 3% BSA in PBS + 0.5% Triton X-100 for 30 min, incubated with primary antibodies overnight at 4 °C, washed with PBS and incubated with secondary antibodies (Thermo Fisher Scientific) plus DAPI (4,6-diamidino2-phenylindole, Sigma) for an additional 2–4hours at room temperature in the dark. Coverslips were washed with PBS and mounted on glass slides using ProLong antifade reagent (Molecular Probes).

For analysis of centrosome numbers immediately after whole genome doubling, the WGD protocol was carried out and mitotic cells were plated onto coverslips in the presence of nocodazole (1μg/ml), reversine (2μM) and BI2536 (100nM) to promote mitotic exit and cytokinesis failure and to allow cells to attach to the coverslip as they entered G1. After 4 h, cells were fixed in PFA and stained as described above. Centrosome numbers were counted by viewing coverslips on a Nikon Ti2-E Eclipse system and a CFI Plan Apochromat λD 40x/0.95 NA air objective (Nikon).

For analysis of p21 levels by cell cycle phase, cells on coverslips were fixed and stained as described above. Coverslips were imaged on a Zeiss Axio Observer using a Plan-apochromat 20×/0.8 M27 Air.

For kinetochore analysis, images were acquired on a Nikon Ti2-E Eclipse system equipped with a heated 37 °C Okolab chamber, using a Kinetix camera (Teledyne Photometrics) with a CFI Plan Apochromat λD 100x/1.45 NA oil objective (Nikon) with NIS-Elements AR software (Nikon). Images were acquired at 1×1 binning and processed using softWorx software, Nikon NIS Elements and ImageJ (National Institutes of Health). Images displayed are maximum intensity projections of deconvolved stacks and were chosen to closely represent the median quantified data. Figure panels were creating using Omero (http://openmicroscopy.org).

The following primary antibodies (all diluted in 3% BSA in PBS) were used at the final concentration indicated: rabbit anti-p21 (#2947, Cell Signaling Technology, 1/800), rabbit anti-53BP1 (Novus biologicals, NB100-304; 1/1000), mouse anti-53BP1 (MAB3802, Millipore, 1/1000), rabbit anti-PLK1 (A300-364A, Bethyl Laboratories, 1/1000), mouse anti-PLK1 (ab17057, Abcam, 1/1000) and guinea pig anti-CENP-C (PD030, Caltag + Medsystems, 1:5000). Secondary antibodies used were highly cross-absorbed goat anti-mouse Alexa Fluor 488 (A-11029), goat anti-mouse Alexa Fluor 568 (A-11031), goat anti-rabbit Alexa Fluor 568 (A-11036) and goat anti-guinea pig Alexa Fluor 647 (A-21450), all used at 1/1000 (Thermo Fisher Scientific). For EdU staining, a base click EdU staining kit was used (BCK-EDU488, Sigma), as per manufacturer’s instructions.

### Quantification of p21 protein levels by cell cycle stage immunofluorescence

Cell cycle segregation of fixed cells was achieved using ImageJ. Firstly, the DAPI channel was used to generate an ROI mask overlay which was then applied to the EdU and p21 channels. Total DAPI signal per mask was determined by measuring mean grey values per ROI in the DAPI channel, background intensity was then subtracted, and the resulting values were multiplied by ROI area to give total DAPI signal per ROI. This total DAPI signal was then plotted against background subtracted EdU intensities from the same ROIs, as seen in figure 4C. From these plots each ROI was gated into G1, S phase, or G2/M and background corrected p21 intensities were plotted for each.

PlotsOfData: https://huygens.science.uva.nl/PlotsOfData was used to generate violin plots. This allows the spread of data to be accurately visualised along with the 95% confidence intervals (thick vertical bars) calculated around the median (thin horizontal lines). This representation allows the statistical comparison between all treatments and timepoints because when the vertical bar of one condition does not overlap with one in another condition the difference between the medians is statistically significant (p < 0.05).

### Kinetochore Image quantification

For quantification of kinetochore protein levels, images of similarly stained experiments were acquired with identical illumination settings and analysed using an ImageJ macro, as described previously^48^. Briefly, the macro performs a threshold and selection of all the kinetochores, using the DAPI and CenpC channels. To generate kinetochore masks, the macro applied a convolution filter to the CenpC channel and perform a threshold selection. The resulting masks were then increased by 1 pixel (to ensure complete kinetochore selection). These masks were used to calculate the relative mean kinetochore intensity 53BP1, PLK1 and CenpC. Fluorescence intensities at kinetochores were normalised to CenpC (i.e. kinetochore marker).

### Time-lapse imaging

For long-term imaging of p53 shRNA RPE1 PLK4-AID cells were first either removed from doxycycline or kept in its presence and incubated for 96hrs (37°C, 5% CO_2_). Cells were then seeded into a Corning Costar 96wp (#3595) at a density of 1000 cells per well either in the presence or absence of doxycycline. Cells were imaged using a Sartorius Incucyte SX5 imaging system and software. Imaging occurred over 7 days with cells being maintained at 37°C and 5% CO_2_ with images being taken using the 10x objective every 4 hours. A total of 3 images were taken of each well per timepoint and mean cell count per well was determined using AI Cell Health analysis.

For brightfield imaging, cells were imaged in a 24-well plate in full growth media in a heated chamber (37°C, 5% CO_2_). For analysis of the mitotic stopwatch, media was supplemented with 4-6μM proTAME (R&D Systems) to promote mild mitotic delays below and above the expected stopwatch threshold of ∼90 min. Only cells entering mitosis in the first 10h were followed for daughter cell fate, and cells which fail to re-enter mitosis within the remaining 186 h window are considered to be arrested.

Images were captured every 4minutes for up to 96 h with a CFI Plan Apochromat λD 10x/0.45 NA air objective (Nikon) using a Nikon Ti2-E Eclipse with a Kinetix camera (Teledyne Photometrics) at 4 × 4 binning or with a 10×/0.5 NA air objective using a Zeiss Axio Observer 7 with a CMOS ORCA flash 4.0 camera at 4 × 4 binning. For analysis of the mitotic stopwatch, only cells entering mitosis in the first 10 h of imaging were followed for daughter cell fate to ensure that daughter cells had sufficient time (>80 h) to enter the next mitosis.

For FUCCI time-lapse imaging, cells were plated at low density (approximately 15,000 cells per well) and imaged in 24-well plates in DMEM inside a heated 37°C chamber with 5% CO_2_. Images were taken every 10 min with a 10×/0.5 NA air objective using a Zeiss Axio Observer 7 with a CMOS ORCA flash 4.0 camera at 4 × 4 binning. Analysis was performed by eye, with 50 random G1 cells being selected for tracking in the first frame of the movie. If a cell exited the field of view during the movie then it was excluded from analysis.

### Western blotting

Protein lysates for western blotting were prepared by scraping cells into Tris-sample buffer (250mM Tris, 10% SDS, 40% Glycerol, 0.1% Bromophenol Blue) or into PBS-sample buffer (4% SDS in PBS, 5% glyverol, 0.1% Bromophenol Blue). Lysates were boiled for 5 min, sonicated (15 s pulse, 50% amp) with a Cole-Palmer Ultrasonic Processor and centrifuged at 13000 rpm for 8 min. Protein concentration was determined by DC assay (BioRad) and 2-mercaptoethanol was added to a final concentration of 5%. Equal amounts of sample were then separated on SDS-PAGE and transferred to 0.45 μm nitrocellulose membranes (Amersham Protran Premium). Membranes were then blocked for 15 min with 5% BSA in TBS with 0.1% Tween 20 (TBS-T) then incubated at 4 °C overnight in 5%BSA TBS-T containing primary antibodies. The following day membranes were washed three times for 5 min each with TBS-T before being transferred to secondary antibodies in a solution of 5% milk in TBS-T. After 1 h of incubation at room temperature membranes were washed three more times in TBS-T and imaged on a LICOR Odyssey CLx system. All blots displayed within each figure panel come from the same experiment and were processed in parallel. The following primary antibodies were used for western blotting: rabbit anti-53BP1 (Novus biologicals, NB100-304; 1/1000), rabbit anti-USP28 (ab126604, Abcam, 1/1000), mouse anti-PLK1 (PLK1 (ab17057, Abcam, 1/1000), mouse anti-MDM2 (33-7100, Thermofisher, 1/1000), mouse anti-p53 (sc-126, Santa Cruz, 1/1000), rabbit anti-p21 (#2947, Cell Signaling Technology, 1/1000), mouse anti-alpha Tubulin (T5168, Sigma, 1/10000), mouse anti-phospho-Histone H2A.X (05-636-I, Millipore, 1/1000) and rabbit anti-alpha Actin (A2066, Sigma, 1/2500). The secondary antibodies were IRDye® 800CW goat anti-mouse (926-32210, LI-COR Biosciences, 1:15000), IRDye® 800CW donkey anti-rabbit (926-32213, LI-COR Biosciences, 1:15000), IRDye® 680RD goat anti-mouse (926-68070, LI-COR Biosciences, 1:15000) and IRDye® 680RD donkey anti-rabbit (926-68071, LI-COR Biosciences, 1:15000). Densitometry was carried out using Image studio lite software V5.2.5 (LI-COR Biosciences).

## References

1. Bielski, C. M. et al. Genome doubling shapes the evolution and prognosis of advanced cancers. Nat Genet 50, 1189–1195 (2018).

2. Zack, T. I. et al. Pan-cancer patterns of somatic copy number alteration. Nat Genet 45, 1134–1140 (2013).

3. Frankell, A. M. et al. The evolution of lung cancer and impact of subclonal selection in TRACERx. Nature 616, 525–533 (2023).

4. Dewhurst, S. M. et al. Tolerance of Whole-Genome Doubling Propagates Chromosomal Instability and Accelerates Cancer Genome Evolution. Cancer Discovery 4, 175–185 (2014).

5. Gerstung, M. et al. The evolutionary history of 2,658 cancers. Nature 578, 122–128 (2020).

6. McPherson, A. et al. Ongoing genome doubling shapes evolvability and immunity in ovarian cancer. Nature 644, 1078–1087 (2025).

7. Vergara, I. A. et al. Evolution of late-stage metastatic melanoma is dominated by aneuploidy and whole genome doubling. Nat Commun 12, 1434 (2021).

8. Storchova, Z. & Pellman, D. From polyploidy to aneuploidy, genome instability and cancer. Nat Rev Mol Cell Biol 5, 45–54 (2004).

9. Aylon, Y. & Oren, M. p53: Guardian of ploidy. Molecular Oncology 5, 315–323 (2011).

10. Ganem, N. J. & Pellman, D. Limiting the Proliferation of Polyploid Cells. Cell 131, 437–440 (2007).

11. Ganem, N. J. et al. Cytokinesis Failure Triggers Hippo Tumor Suppressor Pathway Activation. Cell 158, 833– 848 (2014).

12. Fava, L. L. et al. The PIDDosome activates p53 in response to supernumerary centrosomes. Genes Dev 31, 34–45 (2017).

13. Burigotto, M. et al. Centriolar distal appendages activate the centrosome-PIDDosome-p53 signalling axis via ANKRD26. The EMBO Journal 40, e104844 (2021).

14. Evans, L. T. et al. ANKRD26 recruits PIDD1 to centriolar distal appendages to activate the PIDDosome following centrosome amplification. The EMBO Journal 40, e105106 (2021).

15. Sparr, C. & Meitinger, F. Prolonged mitosis: A key indicator for detecting stressed and damaged cells. Curr Opin Cell Biol 92, 102449 (2025).

16. Sala, C. & Schiebel, E. When the Clock Is Ticking: The Role of Mitotic Duration in Cell Fate Determination. BioEssays n/a, e70061.

17. Lambrus, B. G. et al. p53 protects against genome instability following centriole duplication failure. J Cell Biol 210, 63–77 (2015).

18. Prigent, C. Centriole Duplication at the Crossroads of Cell Cycle Control and Oncogenesis. Cells 14, 1094 (2025).

19. Lambrus, B. G. & Holland, A. J. A New Mode of Mitotic Surveillance. Trends in Cell Biology 27, 314–321 (2017).

20. Uetake, Y. & Sluder, G. Prolonged Prometaphase Blocks Daughter Cell Proliferation Despite Normal Completion of Mitosis. Current Biology 20, 1666–1671 (2010).

21. Meitinger, F. et al. Control of cell proliferation by memories of mitosis. Science 383, 1441–1448 (2024).

22. Lau, T. Y. & Poon, R. Y. C. Whole-Genome Duplication and Genome Instability in Cancer Cells: Double the Trouble. Int J Mol Sci 24, 3733 (2023).

23. Lambrus, B. G. et al. A USP28-53BP1-p53-p21 signaling axis arrests growth after centrosome loss or prolonged mitosis. J Cell Biol 214, 143–153 (2016).

24. Burigotto, M. et al. PLK1 promotes the mitotic surveillance pathway by controlling cytosolic 53BP1 availability. EMBO reports 24, e57234 (2023).

25. Fong, C. S. et al. 53BP1 and USP28 mediate p53-dependent cell cycle arrest in response to centrosome loss and prolonged mitosis. eLife 5, e16270 (2016).

26. Crozier, L. et al. CDK4/6 inhibitor-mediated cell overgrowth triggers osmotic and replication stress to promote senescence. Molecular Cell 83, 4062-4077.e5 (2023).

27. Foy, R. et al. Oncogenic signals prime cancer cells for toxic cell overgrowth during a G1 cell cycle arrest. Molecular Cell 83, 4047-4061.e6 (2023).

28. Manohar, S. et al. Genome homeostasis defects drive enlarged cells into senescence. Molecular Cell 83, 4032-4046.e6 (2023).

29. Crozier, L. et al. CDK4/6 inhibitors induce replication stress to cause long-term cell cycle withdrawal. The EMBO Journal e108599 (2022) doi:10.15252/embj.2021108599.

30. Fulcher, L. J., Sobajima, T., Batley, C., Gibbs-Seymour, I. & Barr, F. A. MDM2 functions as a timer reporting the length of mitosis. Nat Cell Biol 27, 262–272 (2025).

31. Fulcher, L. J., Batley, C., Sobajima, T. & Barr, F. A. Time as a danger signal promoting G1 arrest after mitosis. Trends Cell Biol 0, S0962-8924(25)00126–6 (2025).

32. Yu, Z. K., Gervais, J. L. & Zhang, H. Human CUL-1 associates with the SKP1/SKP2 complex and regulates p21(CIP1/WAF1) and cyclin D proteins. Proc Natl Acad Sci U S A 95, 11324–11329 (1998).

33. Abbas, T. et al. PCNA-dependent regulation of p21 ubiquitylation and degradation via the CRL4Cdt2 ubiquitin ligase complex. Genes Dev 22, 2496–2506 (2008).

34. Wang, W., Nacusi, L., Sheaff, R. J. & Liu, X. Ubiquitination of p21Cip1/WAF1 by SCFSkp2: substrate requirement and ubiquitination site selection. Biochemistry 44, 14553–14564 (2005).

35. Nishitani, H. et al. CDK Inhibitor p21 Is Degraded by a Proliferating Cell Nuclear Antigen-coupled Cul4-DDB1Cdt2 Pathway during S Phase and after UV Irradiation. J Biol Chem 283, 29045–29052 (2008).

36. Manohar, S. & Neurohr, G. E. Too big not to fail: emerging evidence for size-induced senescence. The FEBS Journal (2023) doi:10.1111/febs.16983.

37. Barr, A. R. et al. DNA damage during S-phase mediates the proliferation-quiescence decision in the subsequent G1 via p21 expression. Nature Communications 8, 14728 (2017).

38. Johmura, Y. et al. Necessary and Sufficient Role for a Mitosis Skip in Senescence Induction. Molecular Cell 55, 73–84 (2014).

39. Krenning, L., Feringa, F. M., Shaltiel, I. A., van den Berg, J. & Medema, R. H. Transient Activation of p53 in G2 Phase Is Sufficient to Induce Senescence. Molecular Cell 55, 59–72 (2014).

40. Lanz, M. C. et al. Genome dilution by cell growth drives starvation-like proteome remodeling in mammalian and yeast cells. Nature Structural & Molecular Biology 1–13 (2024) doi:10.1038/s41594-024-01353-z.

41. Gemble, S. et al. Genetic instability from a single S phase after whole-genome duplication. Nature 604, 146– 151 (2022).

42. Vorhauser, J. et al. A redox switch in p21-CDK feedback during G2 phase controls the proliferation-cell cycle exit decision. Mol Cell 85, 3241-3255.e11 (2025).

43. Morrison, L., Loibl, S. & Turner, N. C. The CDK4/6 inhibitor revolution — a game-changing era for breast cancer treatment. Nature Reviews Clinical Oncology 21, 89– 105 (2024).

44. Wander, S. A. et al. The Genomic Landscape of Intrinsic and Acquired Resistance to Cyclin-Dependent Kinase 4/6 Inhibitors in Patients with Hormone Receptor– Positive Metastatic Breast Cancer. Cancer Discovery 10, 1174–1193 (2020).

45. Patnaik, A. et al. Efficacy and Safety of Abemaciclib, an Inhibitor of CDK4 and CDK6, for Patients with Breast Cancer, Non–Small Cell Lung Cancer, and Other Solid Tumors. Cancer Discovery 6, 740–753 (2016).

46. Kudo, R. et al. Long-term breast cancer response to CDK4/6 inhibition defined by TP53-mediated geroconversion. Cancer Cell S153561082400357X (2024) doi:10.1016/j.ccell.2024.09.009.

47. Frank, S. B., Schulz, V. V. & Miranti, C. K. A streamlined method for the design and cloning of shRNAs into an optimized Dox-inducible lentiviral vector. BMC Biotechnol 17, 24 (2017).

48. Saurin, A. T., Waal, M. S. van der, Medema, R. H., Lens, S. & Kops, G. Aurora B potentiates Mps1 activation to ensure rapid checkpoint establishment at the onset of mitosis. Nature Communications 2, 316 (2011).

