## Supplementary Figures for "The mitotic stopwatch limits the proliferation of whole genome doubled cells"

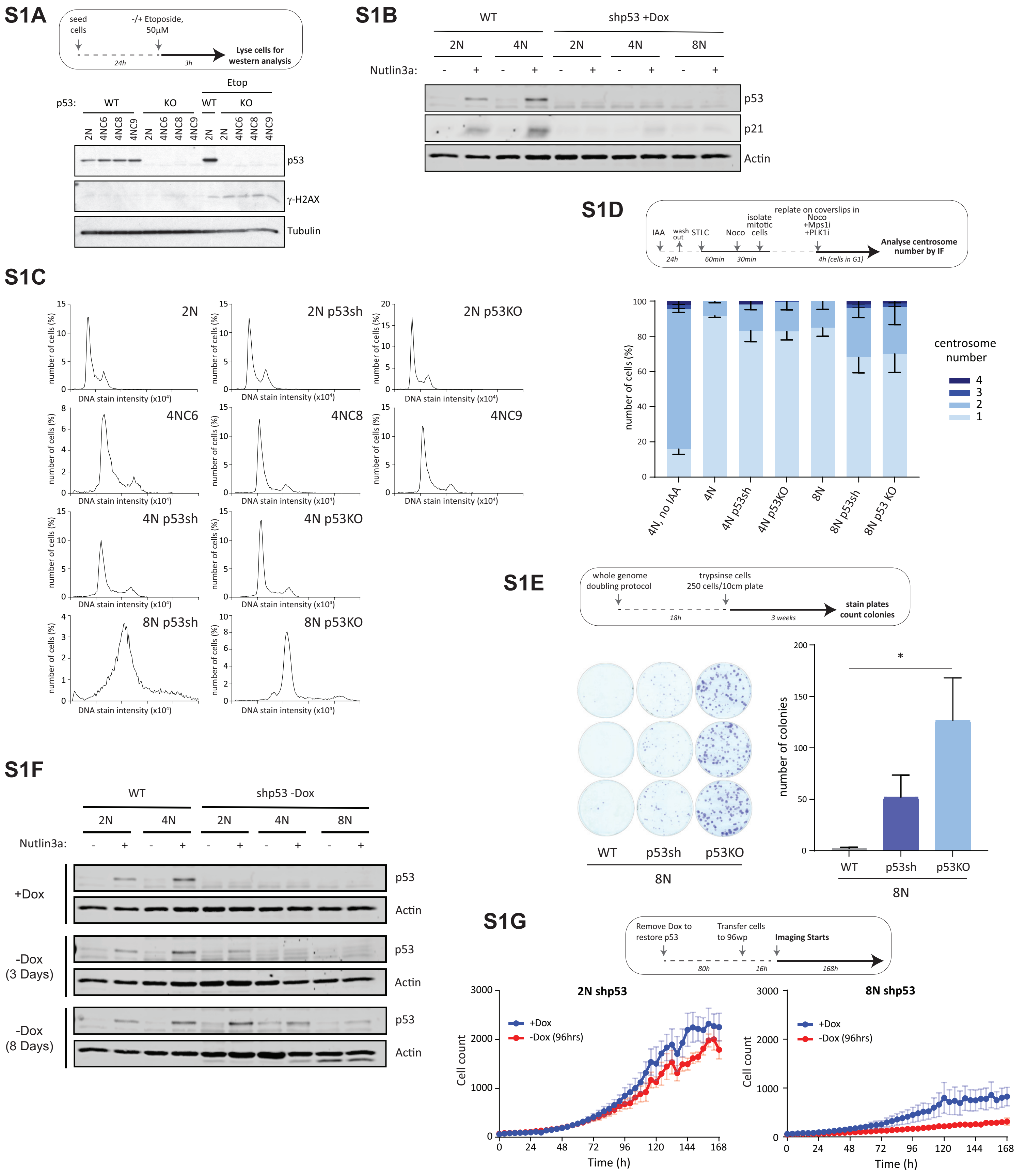


**Supplementary Figure 1** *(related to figure 1)*. **A)** Western blot validating the p53 knockout clones -/+ Etoposide to induce DNA damage ( -H2AX) and stabilise p53. **B)** Western blot validating the conditional knockdown of p53 following treatment with doxycycline in parental (2N), tetraploid (4N), and octoploid (8N) RPE1 PLK4-AID p53 shRNA clones. Cells were treated with/without the MDM2 inhibitor Nutlin3a to stabilize p53. **C)** Propidium Iodide staining of parental (2N), tetraploid (4N), and octoploid (8N) RPE1 PLK4-AID clones with wild-type p53 or following p53 knockout or conditional knockdown with doxycycline inducible p53 shRNA. **D)** Centrosome counts performed immediately following the WGD protocol for the same cell lines shown in S1C. Bars represent mean – SD, n=3. **E)** Colony forming assays of octoploid (8N) cells following the protocol described in 1A. Following the WGD protocol a total of 250 cells were seeded at low density and allowed to form colonies for 3 weeks. Bars represent means ± SD, n = 3. **F)** Western blot validating p53 re-expression in parental (2N), tetraploid (4N), and octoploid (8N) p53 shRNA lines either 3 or 8 days after the removal of doxycycline from cell media. Note that this blot was carried out alongside that shown in S1B and the +Dox condition is the same. **G)** Line graphs representing cell counts of doxycycline-inducible p53-shRNA cells over 7 days either in the presence of doxycycline (blue) or starting 96 hours after doxycycline withdrawal (red). The left graph represents parental (2N) cells while the right graph shows octoploid (8N) cells. Data shown is the mean of 3-5 repeats +/- SEM.

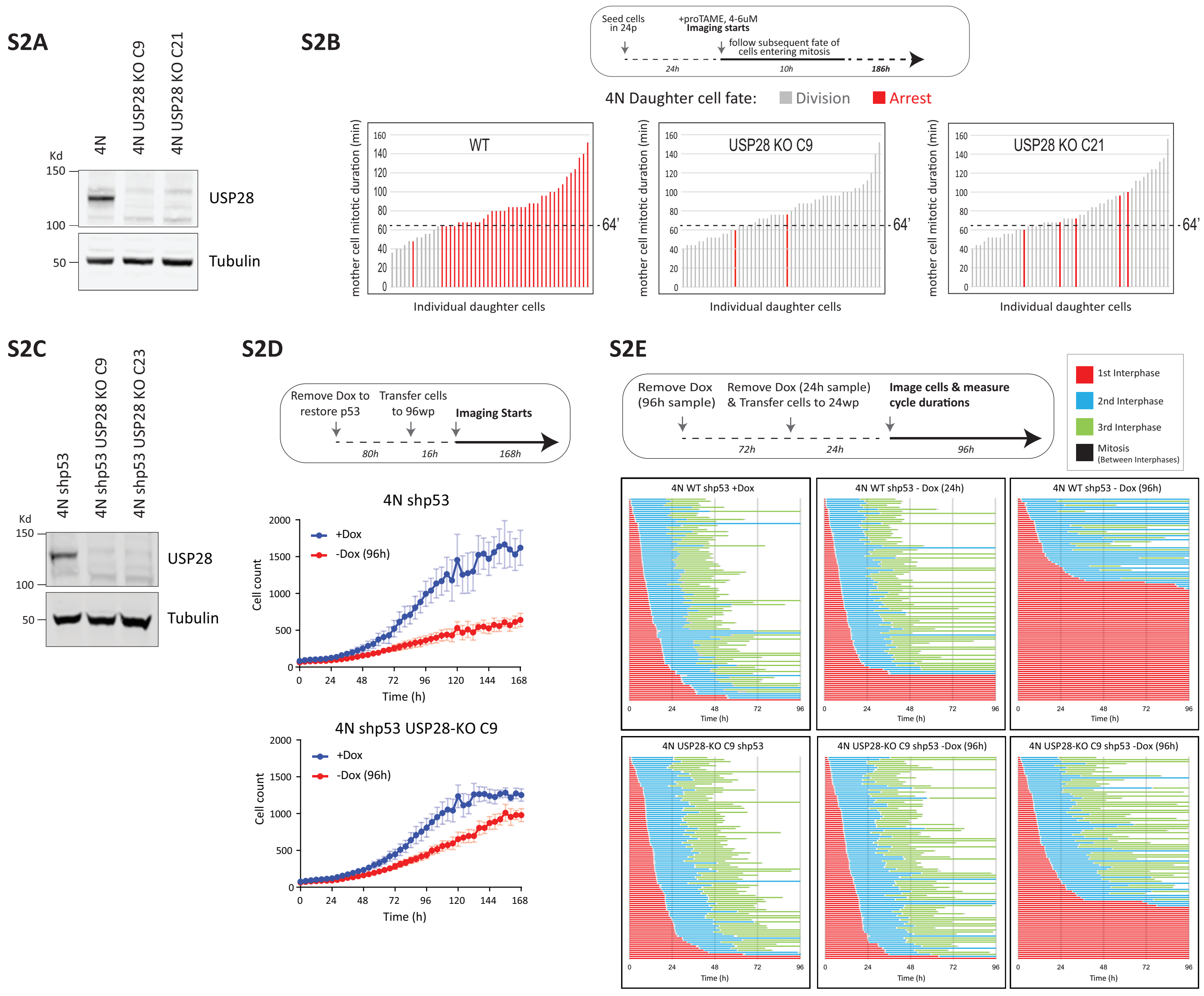


**Supplementary Figure 2** *(related to figure 2)*. **A)** Western blot to validate the knockout of USP28 in lysates from two whole genome doubled (4N) RPE1 PLK4-AID clones. **B)** Plots showing daughter cell fate as a function of mother cell mitotic duration in WT WGD cells and two different USP28-KO WGD clones following treatment with proTAME to induce mitotic delays of varying lengths. Only cells entering mitosis in the first 10h were followed for daughter cell fate. Each bar represents the time spent in mitosis for a single mother cell while the colour of the bar represents daughter cell fate with grey representing continued proliferation and red depicting cell cycle arrest. **C)** Western blot to validate the knockout of USP28 in two whole genome doubled (4N) RPE1 PLK4-AID p53 shRNA clones. **D)** Line graphs representing cell counts of doxycycline-inducible p53-shRNA cells over 7 days either in the presence of doxycycline (blue) or starting 96 hours after doxycycline withdrawal (red). The top graph represents USP28-WT WGD (4N) cells while the bottom graph shows USP238-KO WGD (4N) cells. Data shown is the mean of 3-5 repeats +/- SEM. **E)** Single cell traces of doxycycline-inducible USP28 WT/KO WGD (4N) p53-shRNA cells (each line represents one cell lineage) showing three cell cycles in the presence of doxycycline or starting either 24 or 96 hours after doxycycline withdrawal, as indicated. Fifty cells were analysed at random for each repeat and two experimental repeats are displayed (100 cells total). Note that the graphs for 4N WT shp53 are reproduced from Fig 2D.

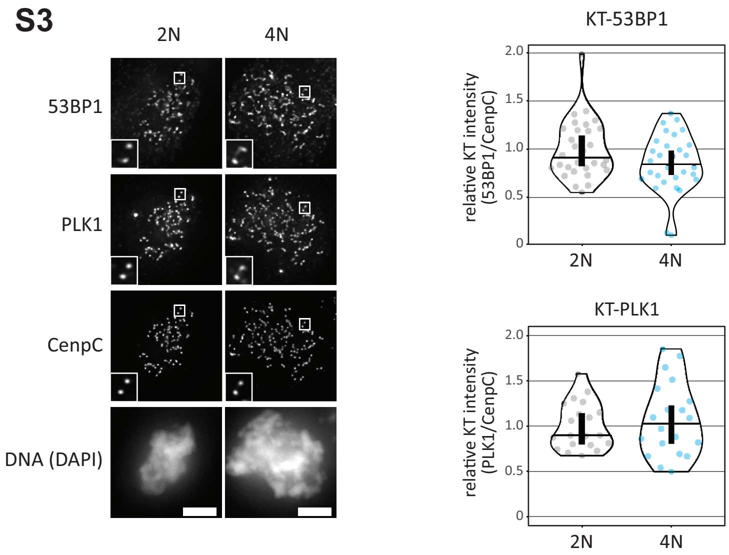


**Supplementary Figure 3** *(related to figure 4)*. Representative immunofluorescence images (left) and quantifications (right) showing 53BP1 and PLK1 at unattached kinetochores in nocodazole-arrested mitotic diploid and tetraploid RPE-PLK4-AID cells, relative to the kinetochore marker, CenpC. The insets show magnifications of the outlined regions. Scale bars: 5 µm. Inset size: 1.5µm. Kinetochore intensities from 20-30 cells, 2 experiments. Violin plots show the distributions of kinetochore intensities between cells. For each violin plot, each dot represents an individual cell, the horizontal line represents the median and the vertical one the 95% CI of the median.
